# A simple offspring-to-mother size ratio predicts post-reproductive lifespan

**DOI:** 10.1101/048835

**Authors:** George Maliha, Coleen T. Murphy

## Abstract

Why do many animals live well beyond their reproductive period? This seems counter to the theory that the fraction of life spent reproducing should be maximized in order to maximize the number of offspring produced in each generation. To resolve this paradox, hypotheses have been developed that evoke parental or grandparental care as reasons for post-reproductive life (e.g., the Mother and Grandmother Hypotheses). However, these hypotheses fail to explain the presence of post-reproductive life in organisms that do not care for their young, such as *Caenorhabditis elegans*. Here we show that a candidate proxy of the stress of childbirth explains a large portion of the variance in post-reproductive lifespans across many species. A remarkably simple metric, the “offspring ratio” (ratio of the size or weight of offspring to that of the mother) explained 77% of the variance of the post-reproductive lifespan in a sample drawn from widely dispersed taxa. Our results suggest that the stress of childbirth is an important and conserved determinant of post-reproductive lifespan. Thus, long post-reproductive lifespan may simply be a byproduct of the somatic health required for reproduction of large progeny, regardless of parental care.

## Introduction

Although post-reproductive life is often thought of as a result of modern medicine’s extensions of lifespan, even before these developments, human females were documented to go through menopause and spend the remainder of their lives without the ability to reproduce (“post-reproductive life”) [1,2]. Women who were able to reproduce late in life (without modern reproductive assistance) also lived longer, suggesting a positive correlation between lifespan and reproductive span [3]. Previous work has hypothesized that while connected, reproductive and total lifespan could be under differential control (perhaps even trading off against one another) [4,5], but the reasons for this differential control have not been elucidated [4,5]. In addition to maternal aging effects on progeny quality, the onset of menopause has serious biological implications due to its effects on normal regulatory processes. For example, rates of cancer, cardiovascular disease, and other degenerative processes drastically increase post-menopause [6-8]. Therefore, elucidating the mechanisms that regulate onset of post-reproductive life and subsequent effects on aging has become more critical.

Several theories have been proposed to explain the existence of post-reproductive life through direct or indirect parental care. For instance, the “Mother Hypothesis” theorizes that females stop reproducing in order to concentrate their efforts and resources in raising already-birthed offspring [9,10]. Moreover, the presence of menopause and post-reproductive life can also protect existing offspring by discouraging males from mating with older mothers [11]. The “Grandmother Hypothesis” posits that post-reproductive females assist in the reproductive success of their daughters, through care of grandprogeny [9,12-14]. Mother and Grandmother Hypotheses concentrate upon direct benefits to children and grandchildren.

However, we and others have shown that *C. elegans*, like women, have a proportionally long post-reproductive life span (PRLS), despite the fact that they do not care for their young [15-19]. In addition, the existence of PRLS has been suggested in studies of brine shrimp (genus *Artermia*), *Drosophila, Mabuya buettneri* (the African Skink, a reptile), and other organisms [20-26]. Mother and Grandmother theories cannot account for the long post-reproductive lives of *C. elegans* and other organisms with non-human social structures. The limiting factor of C. *elegans* reproductive span is oocyte quality decline with age, as it is in mammals [16]. Moreover, oocyte quality is governed by similar gene sets in mammals and worms [16], suggesting that factors that regulate reproductive aging may be conserved evolutionary. Therefore, we wondered whether there is a also conserved determinant of post-reproductive life span (PRLS) across species.

Here we show that such a factor does exist: the ratio of offspring size to mother size correlates well with length of post-reproductive life span across many species. We hypothesize that the offspring ratio may indicate the level of somatic integrity necessary to successfully reproduce, and our tests of this model in *C. elegans* suggest that altering these ratios can have deleterious effects on the mother’s survival during reproduction.

## Methods

### Statistical Analyses

Linear regressions were conducted using native R functionality and the tools available from the “mlbench” package [27,28]. Graphical labels were constructed using the “calibrate” package [29].

### Calculations

Several values were computed from the data: Post-Reproductive Lifespan (“PRLS”), Offspring to Maternal Size Ratio (“Offspring Ratio”), Reproductive Window (the proportion of life in which the species can reproduce), Maturity Proportion (the proportion of life spent maturing to reproductive age), Gestational Proportion (the proportion of life spent in gestation in a single reproductive cycle), and Weaning Proportion (the proportion of life spent weaning offspring from one reproductive cycle; only applicable to mammals).

PRLS was computed as the ratio of life after last documented reproduction to maximum documented life expectancy minus maturity time (the final factor facilitates comparisons outside of mammals) [30]. Offspring ratio was computed by taking the ratio of the offspring to maternal size or weight (depending on the data available). When weights were unavailable, the cube of the length of offspring at birth and mother was used. Litter size-adjusted ratio was computed by multiplying the average size of a litter by the offspring ratio. Reproductive Window was computed as the proportion of maximum lifespan between the age of maturity and reproductive senescence. Maturity Proportion was computed solely for mammals and birds as the ratio of maturation age (for females) to maximum lifespan (the data for non-mammals was not reliable enough). Gestational Proportion was computed similarly as the ratio of gestational time to maximum age.

The equations used to compute the values, then, were as follows:

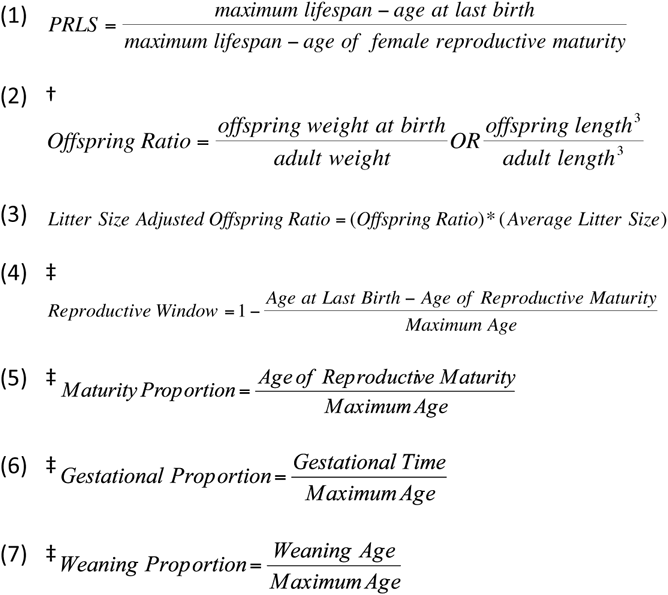

†Offspring Ratio was calculated using weights if possible, but if not, lengths were used. If offspring length could not be found, it was approximated using the length of the female gamete (in the case of sea urchins). While among mammals it has been found that length to the fourth power (not third) relates to body weight, the use of logarithmic regression parameters renders the exact exponent irrelevant. Moreover, as a wide range of animals (not just mammals) were studied, it appeared more appropriate to use the natural geometric relationship [31].

‡This is only computed for mammals and some birds. There was not enough non-mammalian information is available to compute a similar value.

### Data

A dataset was constructed in order to probe various aspects of animal aging. Data for mammals and most birds (both in captivity and in natural habitats) were primarily obtained from anAge: The Animal Aging and Longevity Database and the references contained therein [32]. Reproductive Senescence data was supplemented using previous reviews of primate and mammalian aging [33-35]. The selection of non-mammalian models was guided by previous aging studies [19,36-39]. Data for the following species were also obtained independently: *Strongylocentrotus franciscanus* (Red Sea Urchin) [40-45], *Strongylocentroltus purpuratus* (Purple Sea Urchin) [41-43,46,47], *Oncorhynchus tshawytscha* (Chinook Salmon) [48-51], *Oncorhynchus kisutch* (Coho Salmon) [49,50,52], *Oncorhynchus keta* (Chum Salmon) [42,49,50,53], *Drosophila Melanogaster* [21-24], *Mabuya buettneri* (African Skink, a reptile) [25,26], *Gallus gallus domesticus* (Leghorn-breed chicken) [54-56], *Galeorhinus galeus* (School Shark) [57-60], and *Alligator mississippiensis* (American Alligator) [61-63]. Different measures of longevity (and their values in various species) were obtained from analyses of the ISIS database of Zoo collections [64].

The following assumptions were made (also made explicit in the data tables): In general, the maximum documented age was either obtained from records or computed by summing the longest possible life history for an animal. Age of maturity and reproductive senescence were found by computing averages of given data. Where given, it was favored to take data from the same paper or source in order to maintain consistency among measurements. However, the sources did not fundamentally disagree with one another-and the results were relatively robust to changes in values. For the three species of salmon and two species of sea urchins, a post-reproductive period of one day was assumed (although the assumption was robust to increasing the PRLS and supported by observations), and age to reproductive maturity was assumed to be negligible in the case of the sea urchins because of their relatively long lives [65]. For the shark and alligator, the age of reproductive senescence was assumed to be the average lifespan of the organism, consistent with previous theoretical and empirical work [66]. In addition, age at reproductive maturity was used from either or both genders when considering all mammals-and separated by gender when considering mammals (due to data limitations). Since not explicit, for sea urchins, litter size and birth size was estimated by the number and size of (female) gametes. Since urchins are external fertilizers, though, we assume that this release is the most stressful part of “birthing” and will suffice.

### Data sets

Datatable 1 (Mammals)

Datatable 2 (All Animals)

### Results

To assess whether there is a conserved determinant of post-reproductive life span (PRLS) across species, we gathered information about life history features on a variety of animal taxa (96 species) both within and outside *Mammalia*, including sea urchins, salmon, *Drosophila melanogaster*, and species of reptiles and birds (**Supplemental Table 1**; **Supplemental Table 2**), for which we could obtain information on reproductive senescence, e.g., life span, age at reproductive maturity, adult size, progeny size at birth, and average litter size (see Supplemental information for all available variables). We then compared these features to the post-reproductive life span ratio (Figure 1):

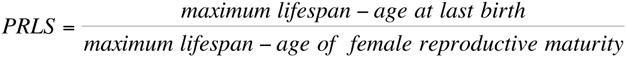

**Figure 1.**
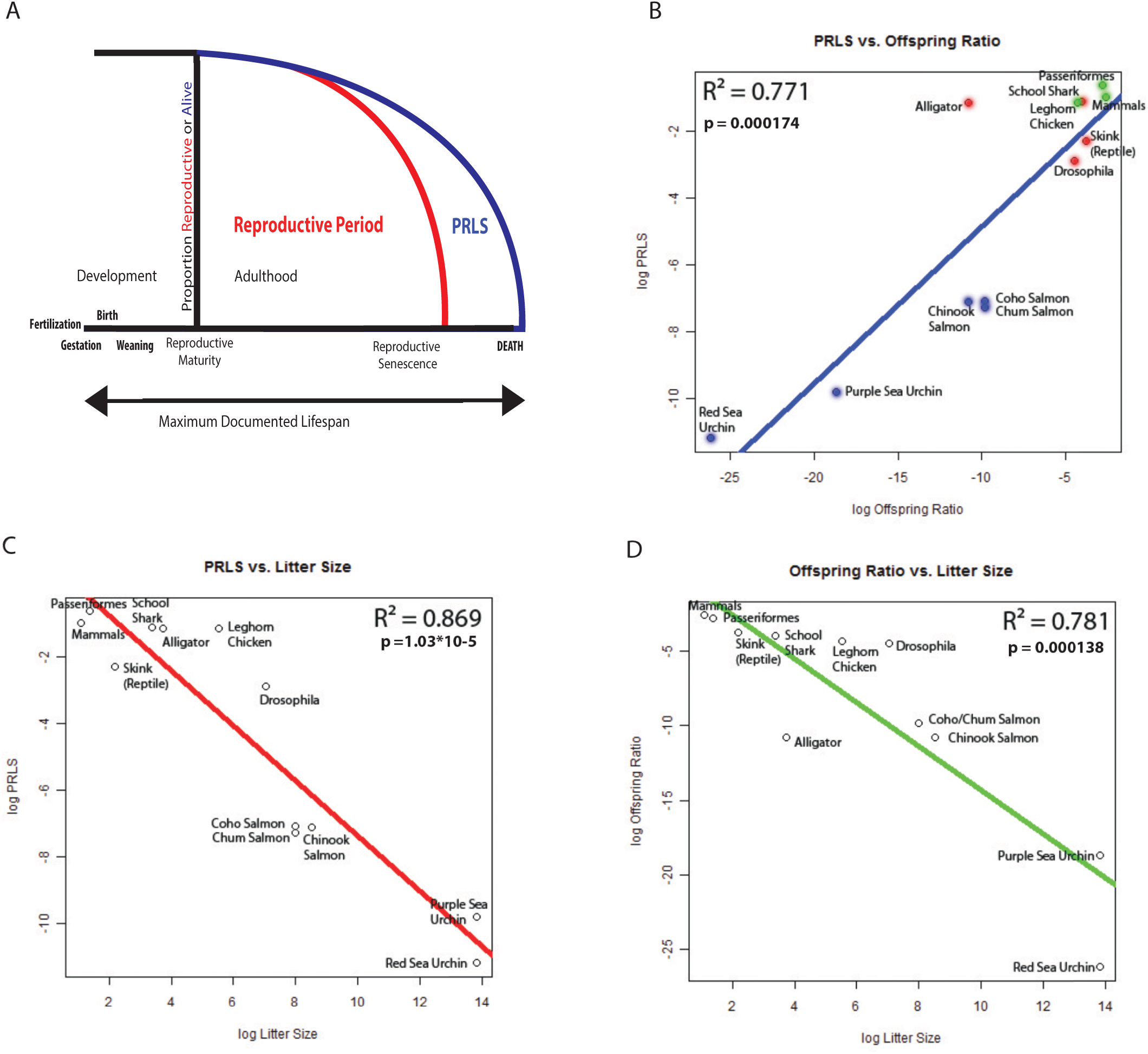
Post-reproductive life span correlates with offspring ratio. A. Scheme of life history parameters considered distinguishing between reproductive and somatic aging. PRLS = “post-reproductive life span” (from reproductive senescence to death). B: Log PRLS is positively correlated with log of the offspring ratio. Regression parameters: Intercept (coefficient estimate: −0.160, standard error: 0.918, t-value: −0.175, and p-value: 0.865) and log offspring ratio: (coefficient estimate: 0.469, standard error: 0.0809, t-value: 5.80, and p-value: 0.000174) on a residual standard error of 1.93, degrees of freedom of 10, R^2^ of 0.771, and an F-statistic of 33.58 on 1 and 10 degrees of freedom. C: log PRLS is negatively correlated with Litter Size, and D: log Offspring Ratio is also negatively correlated with log Litter Size. See **Table 1** for specific regression parameters. All statistical analyses were performed in the R statistical analysis package (64-bit, version 2.14.2). See Supplemental Methods for species information.

**Table 1.** A summary of the regressions relating various parameters to PRLS among a sample of species. Note that litter size and offspring ratio are highly correlated but inversely related.

| Regressions among All Species |  |  |  |  |  |  |  |
| --- | --- | --- | --- | --- | --- | --- | --- |
| y-axis<sup>†</sup> | x-axis<sup>†</sup> | $R^2$ | Coefficient Estimate | Coefficient Standard Error | p-value | Residual Standard Error | Degrees of Freedom |
| PRLS | Litter Size | 0.869 | -0.823 | 0.101 | $1.03 \times 10^{-5}$ | 1.46 | 10 |
| Offspring Ratio | Litter Size | 0.781 | -1.46 | 0.245 | 0.000138 | 3.53 | 10 |
| PRLS | Offspring Ratio | 0.771 | 0.469 | 0.0809 | 0.000174 | 1.93 | 10 |
| PRLS | Litter-Adjusted Offspring Ratio | 0.332 | 0.565 | 0.254 | 0.0500 | 3.29 | 10 |
| PRLS | Maturity Proportion | 0.0846 | 0.932 | 0.969 | 0.359 | 3.85 | 10 |
| <sup>†</sup>Logarithmic transformation of variables unless noted. See supplement for definitions of variables. |  |  |  |  |  |  |  |
| Yellow = t-test of linear regression for selected coefficient is significant at $p < 0.05$ level | | | | | | | |

Within mammals, no one factor accounted for the majority of post-reproductive lifespan, perhaps indicating that several different parameters contribute to the determination of post-reproductive lifespan (**Supplemental Table 1**). In addition, humans (data from the hunter-gatherer-like Hadza and IKung) were not outliers compared to other mammals.

However, a surprisingly simple metric, the ratio of offspring size to mother size, or “Offspring ratio,” correlated well with PRLS across the larger set of species. The measure was computed in two ways depending on data available:

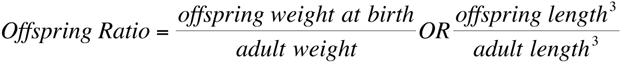

The unadjusted offspring ratio could explain approximately 75% of the variance of post-reproductive lifespan (R^2^ = 0.771; p = 0.000174) (Figure IB). That is, larger offspring with respect to the mother is associated with longer post-reproductive period of the mother. (Note that while litter size has an inverse and highly correlated (R^2^ = 0.869) relationship with PRLS (Figure 1C), it is also correlated with and inversely related to offspring ratio (R^2^ = 0.781), and thus cannot be distinguished from offspring ratio (Figure ID).

In order to attempt to separate the effects of phylogeny from taxonomic adaptations, methods outlined previously were utilized [67,68]. Species-level data among mammals were averaged to the family level, recapitulating similar results and suggesting the presence of adaptation. Graphing the residuals of separate offspring and adult weight regressions against PRLS separated the available placental mammal from the single marsupial data point. As for the non-mammals, in order to capture more data points, we measured offspring ratio through two methods (outlined in Methods). However, the non-mammals are selected from a wide variety of taxa, and even when some of the closely-related species are averaged (e.g., sea urchins and salmon), the results do not change, lessening the concern about these differences emerging from phylogeny (although methods to separate phylogeny and adaptation are unavailable in this case).

Our comparison of offspring ratio to PRLS also roughly groups organisms by reproductive strategy. Clustered in the low offspring ratio/low PRLS area of the graph are red sea urchins, purple sea urchins, and salmon species-all organisms that release unfertilized gametes into the environment (**Figure IB**). The African Skink, *Drosophila melanogaster*, School Shark, and Alligator form another group, producing fertilized embryos that are released into the environment with no parental care (**Figure IB**). Finally, birds and mammals-animals that care for their young-appear to form another cluster at the high offspring ratio/high PRLS region of the graph (**Figure IB**).

When looking at the diversity of animals, it appears that those with the largest offspring ratios have the longest post-reproductive lives, and vice versa. This result is counter to the notion that larger offspring deplete resources, resulting in a shorter post-reproductive life span. Indeed, our results suggest that there is not a direct “tradeoff” between reproductive and post-reproductive life [4]. Instead, we posit that the offspring ratio may be a proxy for stress of childbirth or progeny production. There is likely a point at which an organism cannot devote adequate energy or quality maintenance to reproduction but still has the necessary strength and physical integrity to live. Although in the absence of parental care, neither positive nor negative selection acts on PRLS, these correlations suggest a reason for the particular length of PRLS [5]. In essence, PRLS in many species is simply an unselected residual of life [15,69,70], but the reason for this residual has not been tested previously. We suggest, based on the correlations found here, that more stressful or physically demanding forms of reproduction may require greater strength and integrity in the soma to successfully produce offspring, resulting in greater somatic integrity in the post-reproductive period and correspondingly longer PRLS. The physical manifestation of this threshold can be found when females are pushed to reproduce beyond a typical time: in humans, the most common cause of death of older mothers before the introduction of modern medical interventions was hemorrhaging during childbirth [71]. In the most extreme example in the other direction, sea urchins release millions of one-cell male and female gametes into their environment and reproduce nearly to the end of their ~200 year life span, essentially exhibiting no PRLS.

To test this hypothesis, we perturbed size and reproductive span parameters in *C. elegans*, a model system whose long post-reproductive lifespan has been previously assumed to simply be a lab artifact. [Note: to avoid circularity, we held *C. elegans* out of the correlation analysis in Figure 1.] Our model predicts that if either body size ratio or RS/LS ratio are altered, there would be a suboptimal effect on reproduction or lifespan. TGF-b Sma/Mab mutants are defective in their coupling of longevity and reproductive aging: their germline and oocyte quality is maintained, extending reproductive span (**Figure 2A**), but their somatic tissues age at the same rate as wild-type worms (**Figure 2B**) [15]. Thus, their somatic integrity does not match the high quality of their germlines [16], and their PRLS is compressed without adjusting their offspring ratio proportionally (in fact, their eggs are the same size as wild-type, but their bodies are smaller, thus increasing their offspring ratio) [72]. The effect of this uncoupling is fatal for the animals in late reproduction: because their reproductive span is so long, TGF-b mutants often die from matricide (internal hatching of offspring, **Figure 2C**) while still reproductive, truncating their reproductive lives (**Figure 2D**, * indicates matricide). Wild-type worms usually avoid this fate: *C. elegans* seems to have tied somatic integrity to the offspring ratio, thus tuning post-reproductive lifespan to maximize reproductive span. Further extension of reproductive span without increased somatic integrity results in matricide.

**Figure 2.**
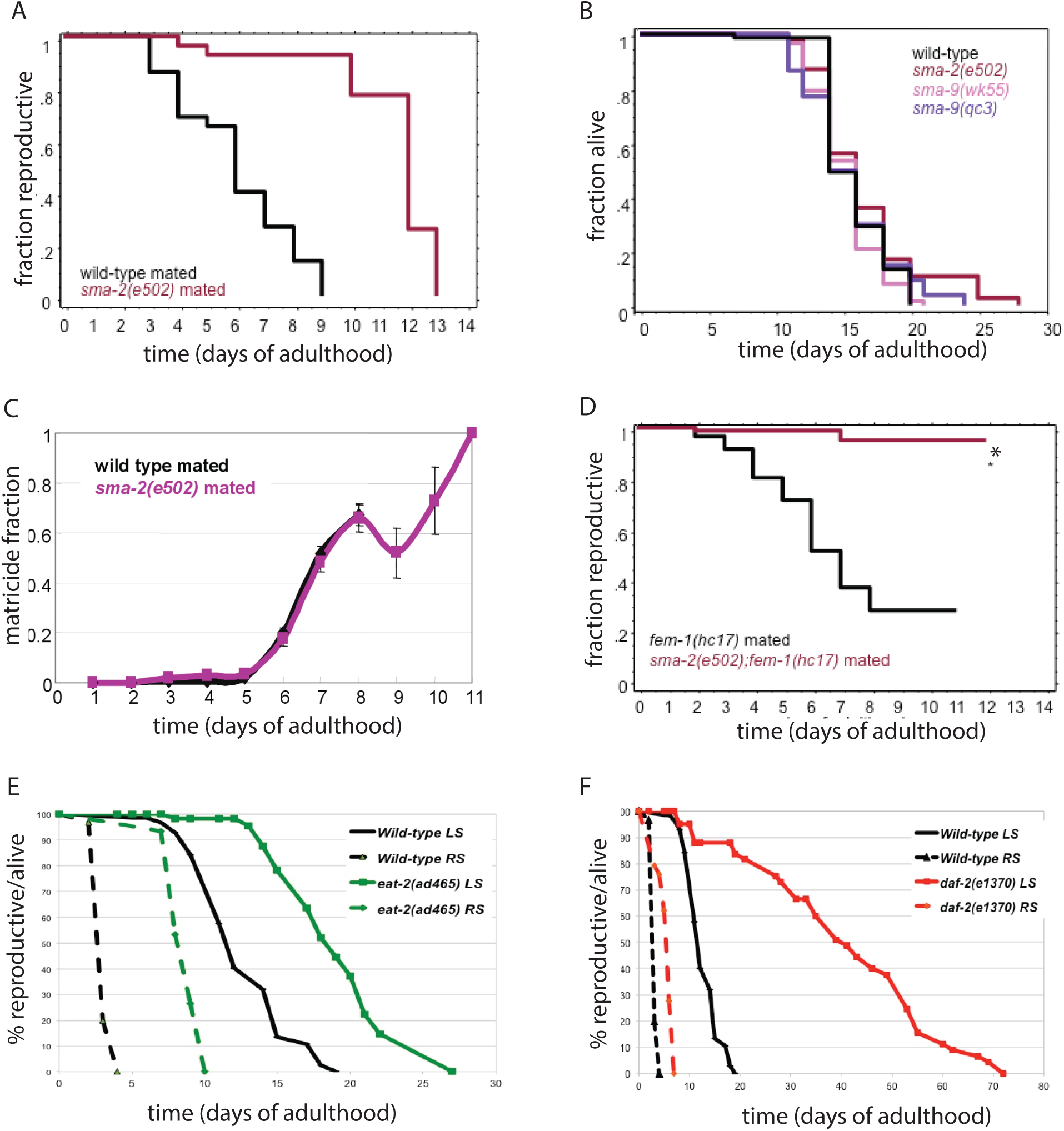
Uncoupling of offspring ratio and reproductive span from longevity results in matricide.A. The reproductive span of the TGF-b mutant *sma-2* is extended relative to wild-type *C. elegans*, but the life span of TGF-b mutants [*sma-2* and *sma-9*) is not extended (**B**). **C**: Matricide rate of TGF-b mutants [*sma-2, sma-9*, and *daf-4*) increases with age. **D**: Matricide (*) causes TGF-b mutant worms to die while still reproductive, [*fem-1* renders the worms spermless, so that all progeny are the result of mating.) E, F. The dietary restriction model *eat-2* and the lnsulin/IGF-1-like receptor mutant *daf-2* both have extended reproductive span (RS) and life span (LS). All figures are adapted from Luo, et al. 2009.

When would the maintenance of somatic integrity extending through late reproduction be important? Limited nutrient conditions, a common situation in the wild, pose such a situation. Dietary restriction extends the lifespan of all animals tested thus far [73]; additionally, dietary restriction delays reproduction [74] (**Figure 2E**). For example, in mammals, fertilized oocyte implantation can be delayed under low nutrient conditions [75], and in humans this process is modulated through FOXO and Insulin signaling [76,77], which also regulates both lifespan and reproductive span in *C. elegans* (**Figure 2F**). In order for reproduction to resume once nutrients become available, the soma must be healthy enough to enable birth, even at advanced ages. Longevity extension, and thus extended PRLS, under dietary-restricted conditions may simply be the result of the coupling of somatic and reproductive aging that is necessary to allow a plastic reproductive response to varying nutrient conditions.

## Discussion

What prevents evolutionary pressure from bringing somatic aging in synchrony with reproductive aging, if for no other reason than to prevent wasting resources that could go to younger generations? For instance, salmon die shortly after spawning; although not instantaneously after reproduction, several days after a multi-year life represents an excellent matching of the two types of aging-despite at least partially separate genetic and regulatory control [69]. Moreover, in the case of salmon specifically, reproduction follows an exhausting migration and severe lack of nutrients (salmon stop feeding), further accelerating somatic decline and death after reproduction [78]. Indeed, lifespan appears to be influenced by hormonal signals (that are in turn partially environmentally influenced) from the reproductive system. For instance, castration of salmon gonads before development prolongs lifespan significantly [79]. Similar findings have been reported in *C. elegans*, mediated through the DAF-12/Nuclear hormone receptor and DAF-16/FOXO signaling pathways [80,81]. Moreover, in humans, the onset of menopause is associated with increased healing time and the rise of cardiovascular disease and other pathologies, and estrogen has been shown to be protective against various health risks-although the mechanisms have not been completely elucidated [82,83]. What remains, though, is to posit why some organisms live very long after they stop reproducing and could theoretically carry another brood. At least a component of this prolonged PRLS could be explained by the nature of biological anti-aging mechanisms. It has been posited that aging is not caused by environmental damage, but rather by the failure to repair that damage [66]. Thus, by that same reasoning, aging depends partly on that damage occurring. If such damage does not occur or occurs more slowly than expected, then aging will be slowed. The case of Werner’s syndrome illustrates; the failure of one such repair mechanism takes years to kill-but eventually it does [84].

On the other hand, in addition to repairing damage to the reproductive tract and even the germ line, reproduction requires “positive control”. That is, certain processes (hormonal, regulatory, or otherwise) must be allowed for reproduction to proceed. If these prerequisites are not met, reproduction halts and the organism enters reproductive senescence. All that has to occur, then, for reproduction to stop-unlike in general somatic aging-is for some prerequisite process to stop or be damaged severely. In a sense, reproduction is much more sensitive to aging because it is a high-stress endeavor and requires so many relatively independent processes at various levels to act. Moreover, these mechanisms are also subject to much greater selection by evolution as they act earlier in life and are intimately concerned with an organism’s evolutionary fitness [85]. Nonetheless, this is not to claim that either type of aging is determined. Medical treatments have intervened in both processes. As witnessed by the dramatic increase in lifespan over the 20th century and the development of in vitro fertilization and general reproductive medicine, both mechanisms of aging can be altered. However, oocyte quality ultimately limits human reproductive span, as in *C. elegans* [69]).

The fact that the simple offspring ratio can correlate parameters across a great number of highly unrelated taxonomic groups, from the most primitive to complex animals, suggests deeper relationships at the genetic and regulatory levels, revealing the intricate connection between reproduction and the structure and parameters of life history. Indeed, previous work has suggested that several factors affect PRLS in less developed organisms in which parental care is a less important factor and social structures differ from those of highly developed mammals [86]. Going forward, it will be essential to develop proxies for predation and parental care [87,88], when appropriate, to account for the remaining variance in PRLS. While parental care may certainly modulate the length of post-reproductive life in some animals, considering offspring size ratio as a proxy for childbirth stress, and PRLS as a byproduct of somatic health during reproduction, offers a new perspective in predicting the post-reproductive lifespan across animal taxa. This model explains the existence of post-reproductive lifespan in animals that do not display parental care, disposing of the need to invoke a “purpose” for PRLS in most species.

## Acknowledgements

We thank Zemer Gitai, Maureen Barr, Daniel Rubenstein, Andrea Bodnar, and members of the Murphy lab for critical discussion of the work, and Shijing Luo and Jasmine Ashraf for assistance with Figure 2.

## Author Contributions

CTM developed the concept; GM carried out all data gathering and analysis; GM and CTM wrote the paper.

## Author Information & Financial Interests Declaration

Original data and sources are available in Supplemental Information. The authors declare no competing financial interests. Correspondence and requests for materials should be addressed to.

## Abbreviations

PRLS: Post-Reproductive Lifespan, RS: Reproductive Span, LS: Life span

---

**Supplemental Table 1.**
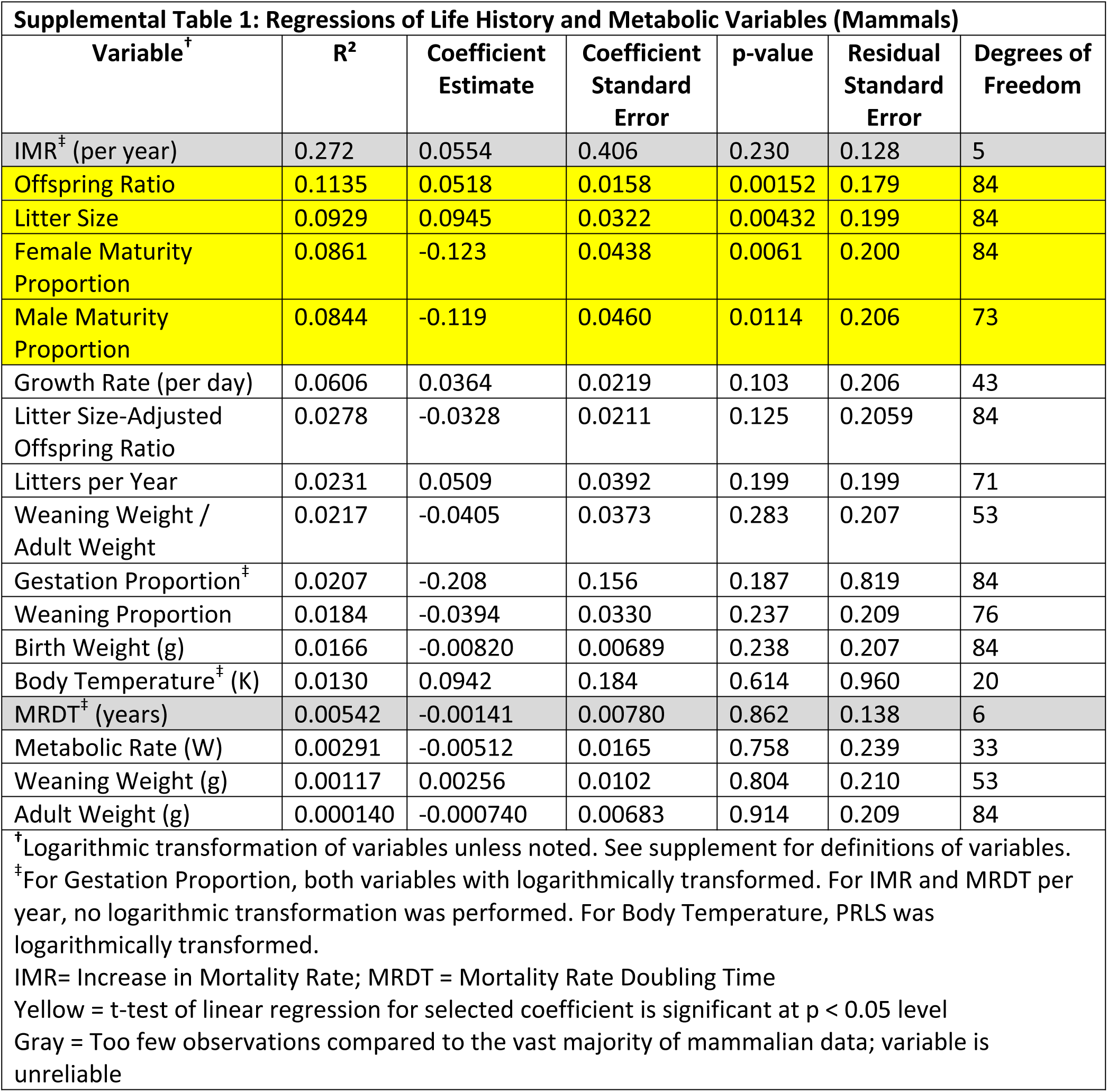
A summary of the regressions performed attempting to relate various parameters to PRLS among a sample of mammals. While other variables could explain around 10% of the variance in PRLS, offspring ratio performed the best (IMR was excluded because there were only 7 data points in the regression versus Offspring Ratio’s 86.). Nonetheless, litter size and the proportion of life spent in “childhood” (for both males and females) also seem to explain a similar proportion of the variance.

| **Supplemental Table 1: Regressions of Life History and Metabolic Variables (Mammals)** |  |  |  |  |  |  |
| --- | --- | --- | --- | --- | --- | --- |
| **Variable<sup>†</sup>** | **R<sup>2</sup>** | **Coefficient Estimate** | **Coefficient Standard Error** | **p-value** | **Residual Standard Error** | **Degrees of Freedom** |
| IMR<sup>‡</sup> (per year) | 0.272 | 0.0554 | 0.406 | 0.230 | 0.128 | 5 |
| Offspring Ratio | 0.1135 | 0.0518 | 0.0158 | 0.00152 | 0.179 | 84 |
| Litter Size | 0.0929 | 0.0945 | 0.0322 | 0.00432 | 0.199 | 84 |
| Female Maturity Proportion | 0.0861 | -0.123 | 0.0438 | 0.0061 | 0.200 | 84 |
| Male Maturity Proportion | 0.0844 | -0.119 | 0.0460 | 0.0114 | 0.206 | 73 |
| Growth Rate (per day) | 0.0606 | 0.0364 | 0.0219 | 0.103 | 0.206 | 43 |
| Litter Size-Adjusted Offspring Ratio | 0.0278 | -0.0328 | 0.0211 | 0.125 | 0.2059 | 84 |
| Litters per Year | 0.0231 | 0.0509 | 0.0392 | 0.199 | 0.199 | 71 |
| Weaning Weight / Adult Weight | 0.0217 | -0.0405 | 0.0373 | 0.283 | 0.207 | 53 |
| Gestation Proportion<sup>‡</sup> | 0.0207 | -0.208 | 0.156 | 0.187 | 0.819 | 84 |
| Weaning Proportion | 0.0184 | -0.0394 | 0.0330 | 0.237 | 0.209 | 76 |
| Birth Weight (g) | 0.0166 | -0.00820 | 0.00689 | 0.238 | 0.207 | 84 |
| Body Temperature<sup>‡</sup> (K) | 0.0130 | 0.0942 | 0.184 | 0.614 | 0.960 | 20 |
| MRDT<sup>‡</sup> (years) | 0.00542 | -0.00141 | 0.00780 | 0.862 | 0.138 | 6 |
| Metabolic Rate (W) | 0.00291 | -0.00512 | 0.0165 | 0.758 | 0.239 | 33 |
| Weaning Weight (g) | 0.00117 | 0.00256 | 0.0102 | 0.804 | 0.210 | 53 |
| Adult Weight (g) | 0.000140 | -0.000740 | 0.00683 | 0.914 | 0.209 | 84 |
| <sup>†</sup>Logarithmic transformation of variables unless noted. See supplement for definitions of variables.<br><sup>‡</sup>For Gestation Proportion, both variables with logarithmically transformed. For IMR and MRDT per year, no logarithmic transformation was performed. For Body Temperature, PRLS was logarithmically transformed.<br>IMR= Increase in Mortality Rate; MRDT = Mortality Rate Doubling Time<br>Yellow = t-test of linear regression for selected coefficient is significant at p < 0.05 level<br>Gray = Too few observations compared to the vast majority of mammalian data; variable is unreliable |  |  |  |  |  |  |

**Supplemental Table 2.**
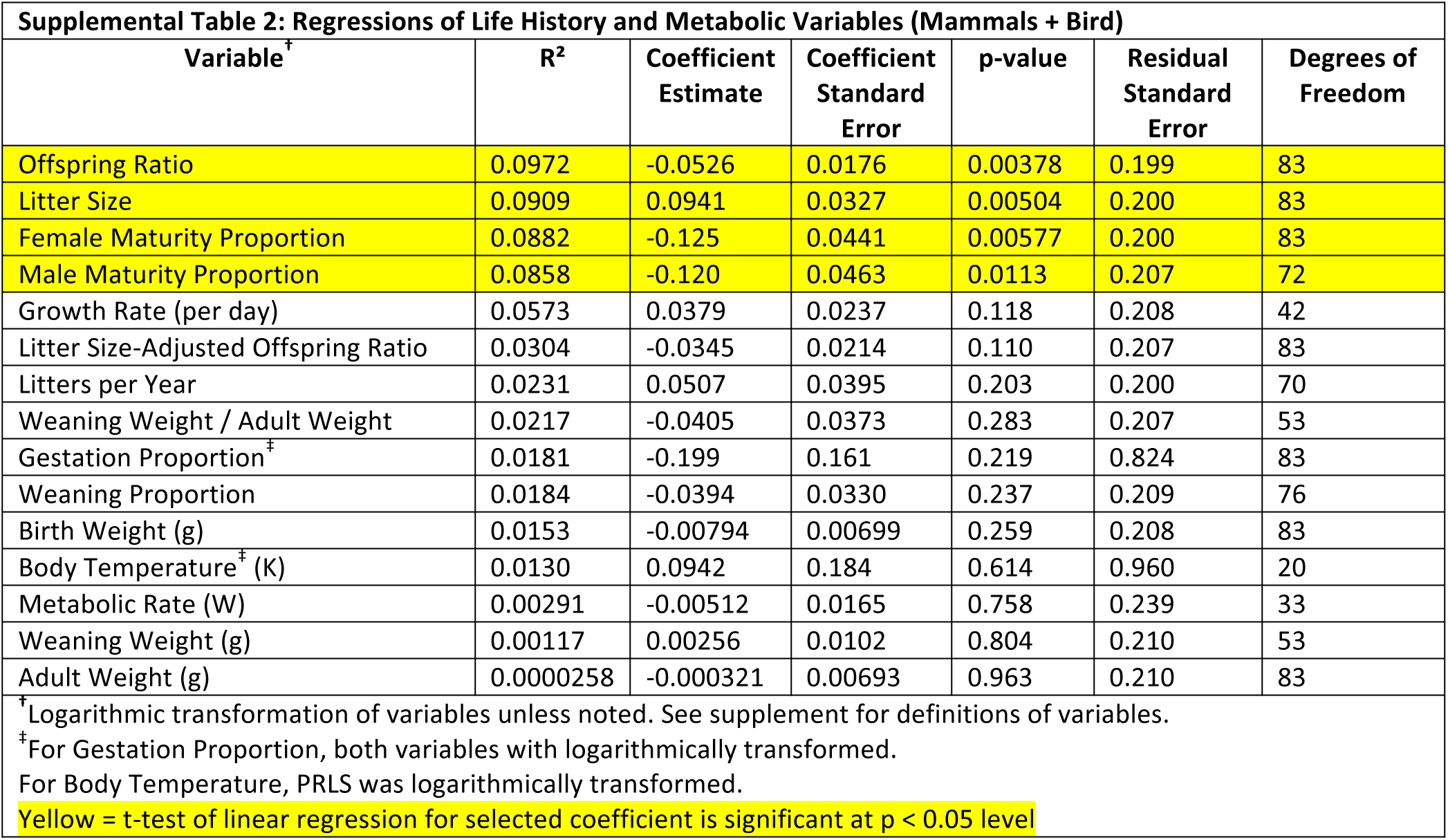
Summary of the regressions relating various parameters to PRLS among a sample of mammals and a bird, sorted by R^2^ value. While other variables could explain around 10% of the variance in PRLS, offspring ratio performed the best. Litter size and the proportion of life spent in “childhood” (for both males and females) explain a similar proportion of the variance. Too few observations of Mortality Rate Doubling time (7) and Increase in Mortality Rate (6) were available to reach significance.

**Supplemental Table 3.**
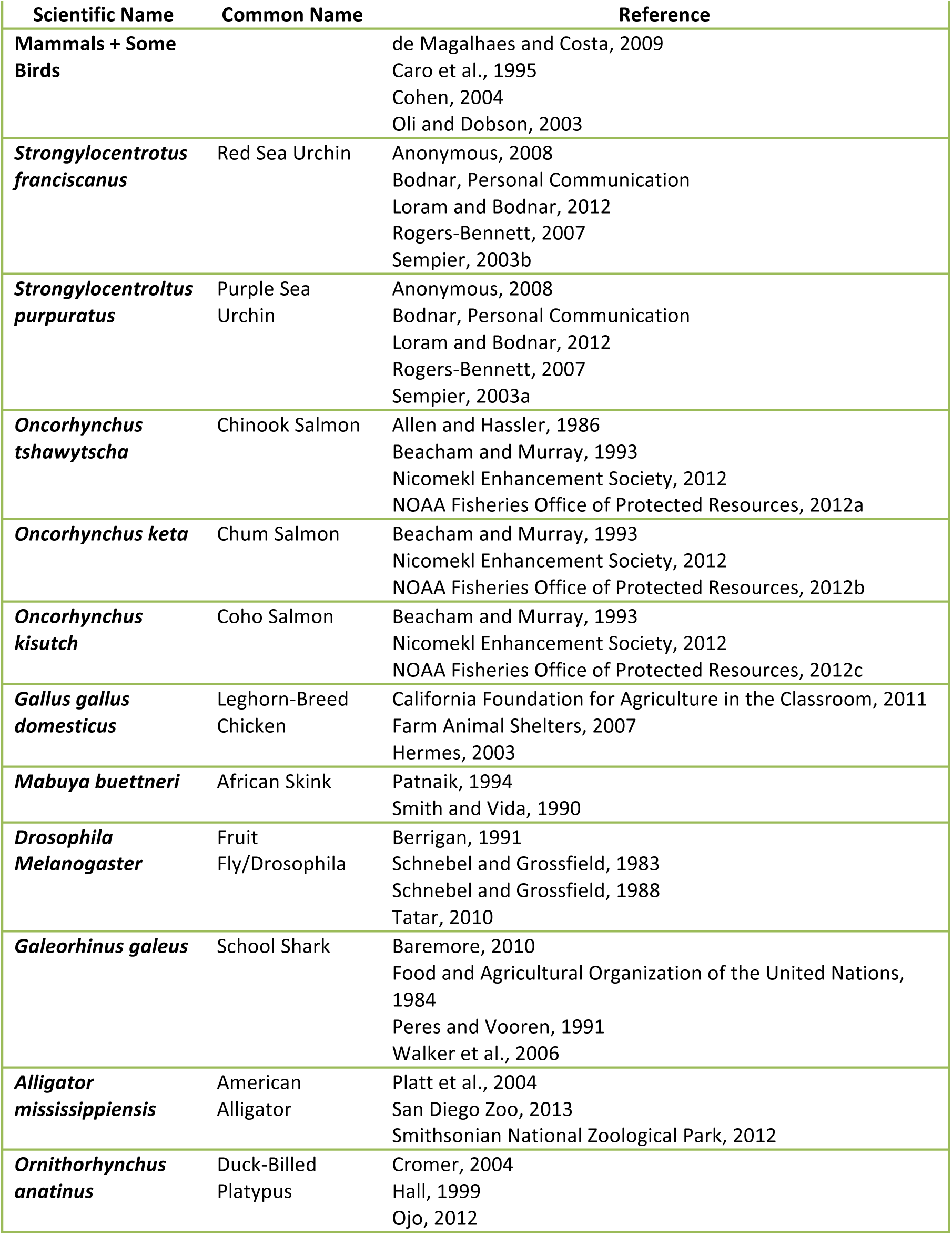
Summary of data sources.

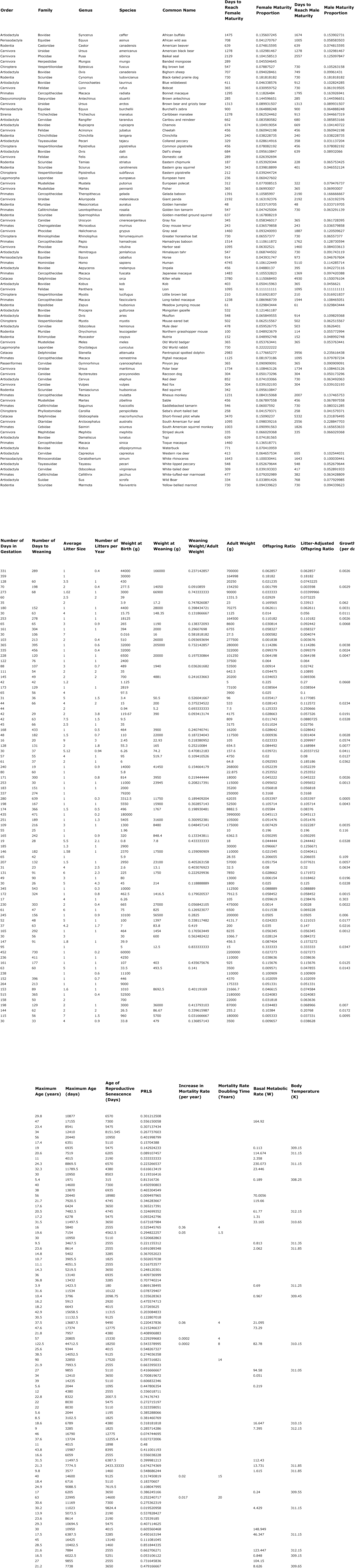
Datatable 1

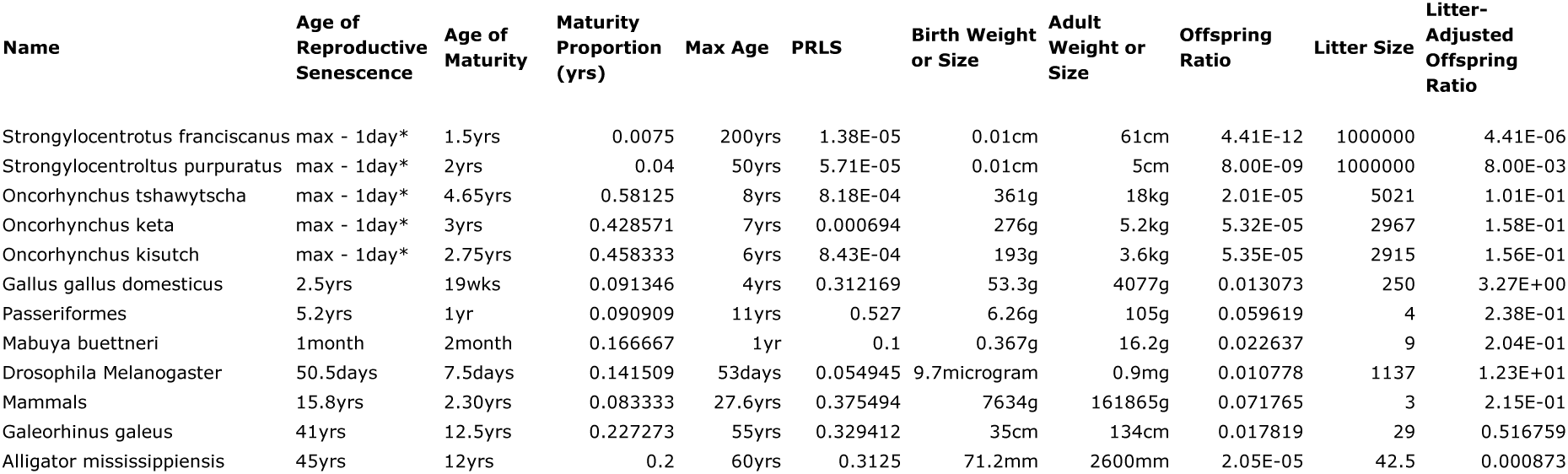
Datatable 2

**Supplemental Figure 1.**
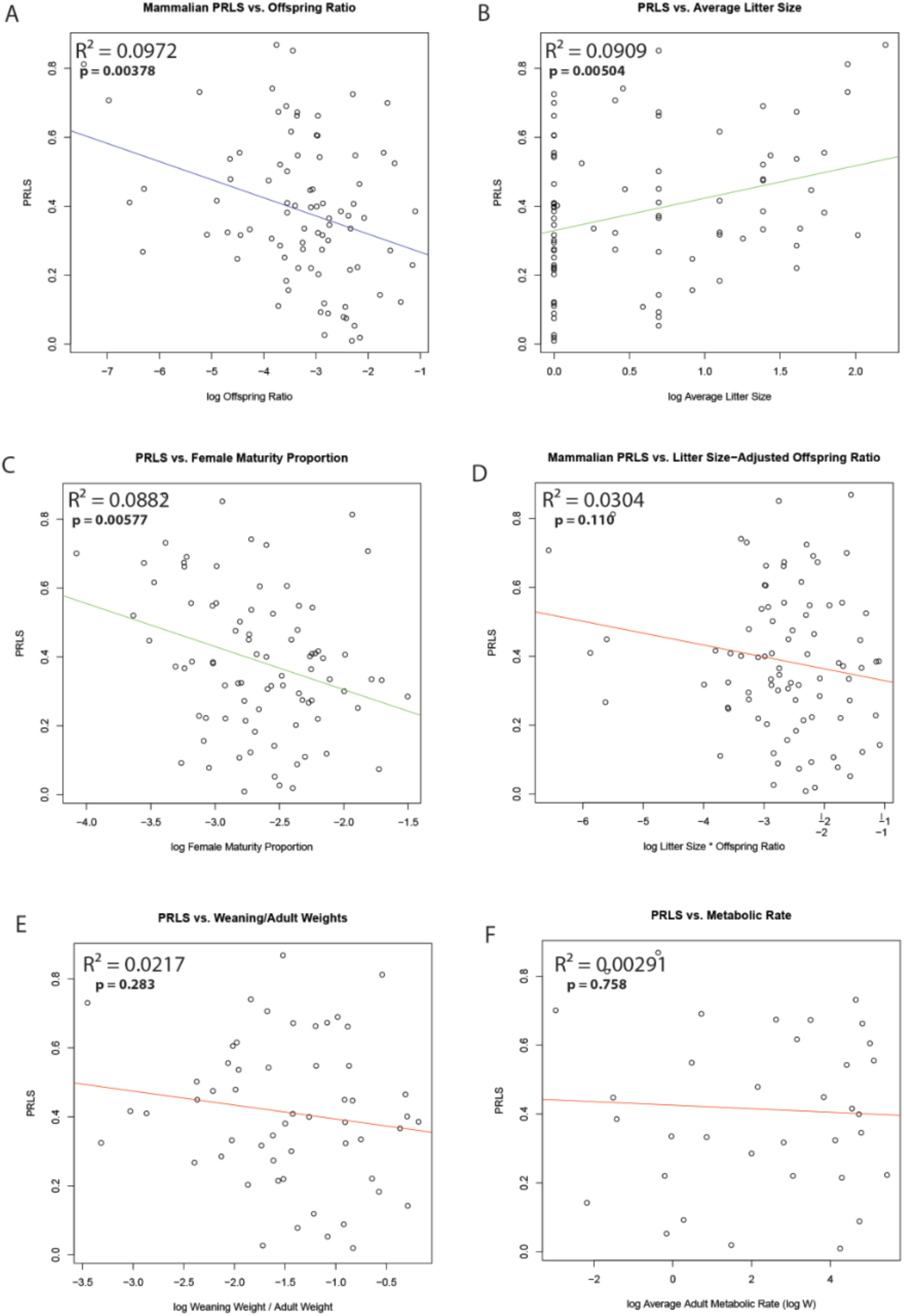
Linear regression of PRLS vs. various variables for 85 mammals distributed across various families and orders. Linear regressions were constructed based on the logarithms of the variable offspring ratio. A: PRLS is positively correlated with log of offspring ratio. Regression parameters: Intercept (coefficient estimate: 0.217, standard error: 0.0613, t-value: 3.54, and p-value: < 0.000659) and log offspring ratio: (coefficient estimate: −0.0522, standard error: 0.0175, t-value: −2.98, and p-value: 0.00378) on a residual standard error of 0.200, degrees of freedom of 83, R^2^ of 0.0956, and an F-statistic of 8.88 on 1 and 83 degrees of freedom. B, C: a sampling of regressions of PRLS against other statistically significant variables: average litter size and female maturity proportion. D-F: a sampling of regressions of PRLS against other statistically non-significant variables: litter size-adjusted offspring ratio, weaning/adult weights, and metabolic rate, in that order. See **Datatable 1** for specific regression parameters. Mammals from the following families are represented (see supplement for detailed information on species): *Bovidae, Cervidae, Suidae, Tayassuidae, Canidae, Felidae, Herpestidae, Mephitidae, Otariidae, Phocidae, Ursidae, Delphinidae, Phyllostomidae, Rhinolophidae, Vespertilionidae, Dasyuridae, Leporidae, Equidae, Rhinocerotidae, Callitrichidae, Cebidae, Cercopithecidae, Hominidae, Castoridae, Chinchillidae, Dipodidae, Echimyidae, Muridae, Sciuridae*, and *Trichechidae*. All statistical analyses were performed in the R statistical analysis package (64-bit, version 2.14.2).

**Supplemental Figure 2.**
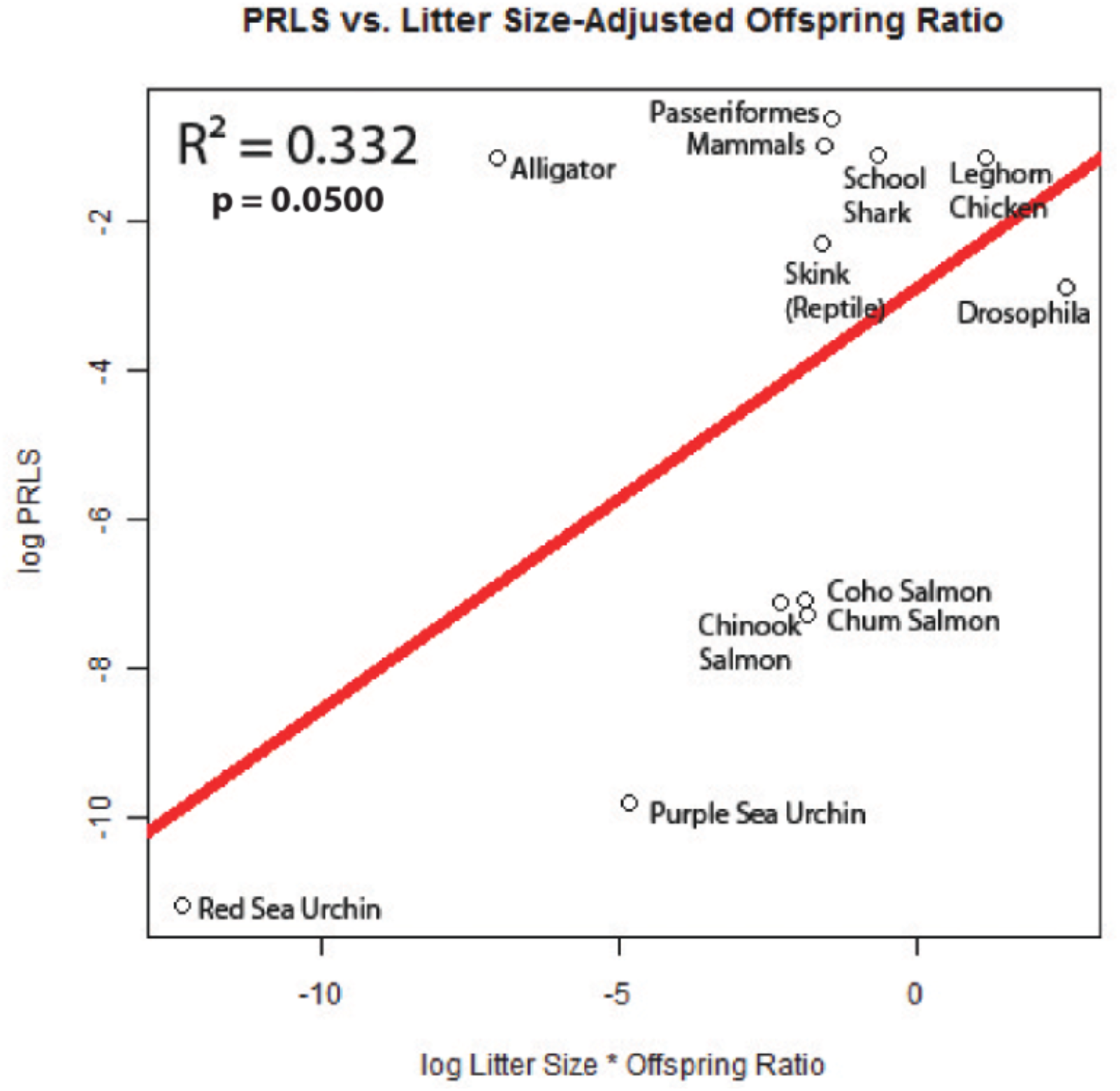
The linear regression of PRLS with litter size-adjusted offspring ratio for the all-animal sample. While the general trend remains (a positive correlation), statistical significance and the ability to explain the variance of the data (R^2^) are less than without litter size. Regression of PRLS against offspring ratio and litter size yield similar results. All statistical analyses were performed in the R statistical analysis package (64-bit, version 2.14.2).

**Supplemental Figure 3.**
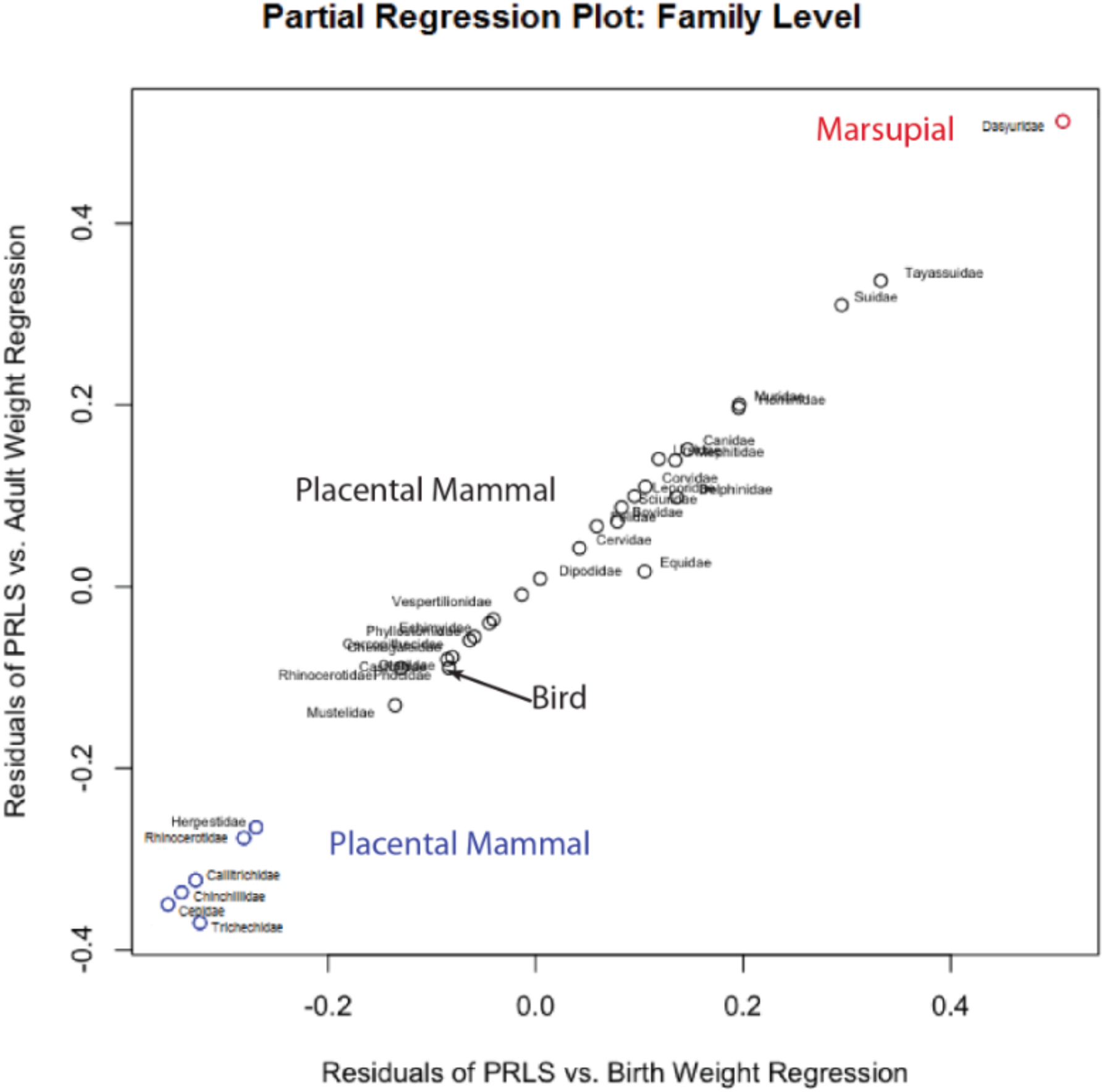
The Partial Regression Plot of the residuals of PRLS vs. Birth Weight Regression and PRLS vs. Adult Weight Regression for 31 families of placental mammals, 1 family of marsupials, and 1 family of birds. The residuals of the separate regressions (PRLS vs. Birth Weight and PRLS vs. Adult Weight) were plotted against each other to extract any natural groupings with respect to PRLS. While the bird family (Corvidae) was indistinguishable from the placental mammals, the marsupial family (Dasyuridae) marked in red readily separated from the placental in the top right hand corner. In addition, a group of diverse families (marked in blue), including Manatees (Trichechidae), Chinchillas (Chinchillidae), and New World Monkeys, appear to separate out in the bottom left hand corner. Key: see Data Table 1 (the families represented are stated in Supplemental Figure 1, with the addition of *Corvidae)*.

## References

1. Caspari R, Lee S. (2004) Older age becomes common late in human evolution. Proceedings of the National Academy of Sciences of the United States of America 101: 10895–900.

2. Amundsen DW, Diers CJ. (1973) The age of menopause in medieval europe. Human Biology 45: 605–12.

3. Perls TT, Alpert L, Fretts RC. (1997) Middle-aged mothers live longer. Nature 389: 133–133.

4. Cohen AA. (2004) Female post-reproductive lifespan: A general mammalian trait. Biological Reviews of The Cambridge Philosophical Society 79: 733–50.

5. Kirkwood TBL, Shanley DP. (2010) The connections between gerneral and reproductive senescence and the evolutionary basis of menopause. Annals of the New York Academy of Sciences 1204: 21–9.

6. Roger V, Go AS, Lloyd-Jones D, Benjamin EJ, Berry JD, et al. (2012) Heart disease and stroke statistics-2012 update: A report from the american heart association. Circulation 125: e2–e220.

7. Howlader N, Noone AM, Krapcho M, Neyman N, Aminou R, et al. (2012) SEER cancer statistics review, 1975–2009. Bethesda, MD: National Cancer Institute.

8. Maas AHEM, Franke HR. (2009) Women’s health in menopause with a focus on hypertension. Netherlands Heart Journal 17: 68–72.

9. Wu JM, Zelinski MB, Ingram DK, Ottinger MA. (2005) Ovarian aging and menopause: Current theories, hypotheses, and research models. Experimental Biology and Medicine 230: 818–28.

10. Madrigal L, Melendez-Obando M. (2008) Grandmothers’ longevity negatively affects daughters’ fertility. American Journal of Physical Anthropology 136: 223–9.

11. Turke PW. (1997) Hypothesis: Menopause discourages infanticide and encourages continued investment by agnates. Evolution and Human Behavior 18: 3–13.

12. Hawkes K. (2004) The grandmother effect. Nature 428: 128–9.

13. Hawkes K. (2003) Grandmothers and the evolution of human longevity. American Journal of Human Biology 15: 380–400.

14. Hawkes K, O’Connell JF, Jones NG, Alvarez H, Charnov EL. (1998) Grandmothering, menopause, and the evolution of human life histories. Proceedings of the National Academy of Sciences of the United States of America 95: 1336–9.

15. Luo S, Shaw WM, Ashraf J, Murphy CT. (2009) TGF-beta sma/mab signaling mutations uncouple reproductive aging from somatic aging. PLoS Genetics 5: el000789.

16. Luo S, Kleemann GA, Ashraf JM, Shaw WM, Murphy CT. (2010) TGF-B and insulin signaling regulate reproductive aging via oocyte and germline quality maintenance. Cell 15: 299–312.

17. Hughes SE, Evason K, Xiong C, Kornfeld K. (2007) Genetic and pharmacological factors that influence reproductive aging in nematodes. PLoS genetics 3: e25.

18. Huang C, Xiong C, Kornfeld K. (2004) Measurements of age-related changes of physiological processes that predict lifespan of caenorhabditis elegans. Proceedings of the National Academy of Sciences of the United States of America 101: 8084–9.

19. Austad S, Hazzard DG, Warner HR, Finch CE. (1991) National institution on aging, NIH, workshop on alternative animal models for research on aging: Invertebrates. Experimental Gerontology 26: 432–8.

20. Agh N, Van Stappen G, Bossier P, Sepehri H, Lofti V, et al. (2008) Effects of salinity on survival, growth, reproductive and life span characteristics of artemia populations from urmia lake and neighboring lagoons. Pakistan Journal of Biological Sciences 11: 164–72.

21. Tatar M. (2010) Reproductive aging in invertebrate genetic models. Annals of the New York Academy of Sciences 1204: 149–55.

22. Berrigan D. (1991) The allometry of egg size and number in insects. Oikos 60: 313–21.

23. Schnebel EM, Grossfield J. (1983) A comparison of lifespan characteristics mDrosophila. Experimental Gerontology 18: 325–37.

24. Schnebel EM, Grossfield J. (1988) Antagonistic pleiotropy: An interspecific drosophila comparison. Evolution 42: 306–11.

25. Smith JM, Vida G. (1990) Organizational constraints on the dynamics of evolution. New York, New York: Manchester University Press.

26. Patnaik BK. (1994) Aging in reptiles. Gerontology 40: 200–20.

27. R Development Core Team. (2012) R: A language and environment for statistical computing. Vienna, Austria: R Foundation for Statistical Computing.

28. Leisch F, Dimitriadou E. (2010) Mlbench: Machine learning benchmark problems.: R package version 2. 1–1.

29. Graffelman J. (2012) Calibrate: Calibration of scatterplot and biplot axes.

30. Liu G, Rogers J, Murphy CT, Rongo C. (2011) EGF signaling activates the ubiquitin proteasome system to modulate C. elegans lifespan. EMBO Journal 30: 2990–3003.

31. May RM, Rubenstein Dl. (1986) Reproductive strategies. In: Austin CR, Short RV, editors. Reproduction in Mammals. Cambridge, UK: Cambridge University Press. pp. 1–23.

32. de Magalhaes JP, Costa J. (2009) A database of vertebrate longevity records and their relation to other life-history traits. Journal of Evolutionary Biology 22: 1770–4.

33. Cohen AA. (2004) Female post-reproductive lifespan: A general mammalian trait. Biological Reviews of the Cambridge Philosophical Society 79: 733–50.

34. Cam TM, Sellen DW, Parish A, Frank R, Brown DM, et al. (1995) Termination of reproduction in nonhuman and human female primates. International Journal of Primatology 16: 205–20.

35. Oli MK, Dobson FS. (2003) The relative importance of life-history variables to population growth rate in mammals: Cole’s prediction revisited. American Naturalist 161: 422–40.

36. Hazzard DG, Warner HR, Finch CE. (1991) National institution on aging, NIH, workshop on alternative animal models for research on aging: Introduction. Experimental Gerontology 26: 411–12.

37. Cheal M, Hazzard DG, Warner HR, Finch CE. (1991) National institution on aging, NIH, workshop on alternative animal models for research on aging: Mammals. Experimental Gerontology 26: 412–7.

38. Ottinger MA, Hazzard DG, Warner HR, Finch CE. (1991) National institution on aging, NIH, workshop on alternative animal models for research on aging: Birds. Experimental Gerontology 26: 426–32.

39. Schreibman M, Hazzard DG, Warner HR, Finch CE. (1991) National institution on aging, NIH, workshop on alternative animal models for research on aging: Reptiles and fish. Experimental Gerontology 26: 417–26.

40. Sempier S. (2003) Red sea urchmStrongylocentrotus franciscanus. 2012.

41. [Anonymous]. (2008) Sea urchin embryology: Gametes. 2012.

42. Loram J, Bodnar A. (2012) Age-related changes in gene expression tissues of the sea urchmStrongylocentrotus purpuratus. Mechanisms of Ageing and Development 2012: 338–47.

43. Rogers-Bennett L. (2007) The ecology ofStrongylocentrotus franciscanusandStrongylocentrotus purpuratus. In: Lawrence J, editor. Sea Urchins: Biology and Ecology. Amsterdam, Netherlands: Elsevier, pp. 398.

44. Ebert T. (2008) Longevity and lack of senescence in the red sea urchinStrongylocentratus franciscanus. Experimental Gerontology 43: 734–8.

45. Ebert T, Southon J. (2003) Red sea urchins (Strongylocentrotus franciscanus) can live over 100 years: Confirmation with A-bomb^14^carbon. Fishery Bulletin, United States 101: 915–22.

46. Sempier S. (2003) Purple sea urch\n:Strongylocentrotus purpuratus. 2012.

47. Ebert T. (1967) Negative growth and longevity in the purple sea urchin strongylocentrotus purpuratus (stimpson). Science 157: 557–8.

48. Allen MA, Hassler TJ. (1986) Species profiles: Life histories and environmental requirements of costal fishes and invertebrates (pacific southwest)-chinook salmon. U S Fish Wildlife Service Biological Reports: 26.

49. Beacham TD, Murray CB. (1993) Fecundity and egg size variation in north american pacific salmon (Oncorhynchus). Journal of Fish Biology 42: 485–508.

50. Nicomekl Enhancement Society. (2012) Our salmon species/life cycle. 2012.

51. NOAA Fisheries Office of Protected Resources. (2012) Chinook salmon (Oncorhynchus tshawytscha). 2012.

52. NOAA Fisheries Office of Protected Resources. (2012) Coho salmon (Oncorhynchus kisutch). 2012.

53. NOAA Fisheries Office of Protected Resources. (2012) Chum salmon (Oncorhynchus keta). 2012.

54. California Foundation for Agriculture in the Classroom. (2011) Commodity fact sheet: Eggs. 2012.

55. Farm Animal Shelters. (2007) Farm animal care: Chicken care. 2012.

56. Hermes JC. (2003) Why did my chickens stop laying? PNW 565: 1–2.

57. Peres MB, Vooren CM. (1991) Sexual development, reproductive cycle, and fecundity of the school SharkGaleorhinus galeusoff southern brazil. Fishery Bulletin, United States 89: 655–67.

58. Baremore IE. (2010) Reproductive aspects of the atlantic angel sharkSquatina dumeril. Journal of Fish Biology 76: 1682–95.

59. Walker Tl, Cavanagh RD, Stevens JD, Carlisle AB, Chiaramonte GE, et al. (2006) Galeorhinus galeus. 2013.

60. Food and Agricultural Organization of the United Nations. (1984) Species fact sheets\Galeorhinusgaleus. 2013.

61. Piatt SG, Resetar A, Stuart B. L. (2004) Maximum clutch size of the american alligator. Florida Field Naturalist 32: 102–6.

62. San Diego Zoo. (2013) San diego zoo’s animal bytes: Alligator & crocodile. 2013.

63. Smithsonian National Zoological Park. (2012) Fact sheets: American alligator. 2013.

64. Moorad JA, Promislow DEL, Fiesness N, Miller RA. (2012) A comparative assessment of univariate longevity measures using zoological animals records. Aging Cell 11: 940–9.

65. Shapovalov L, Taft A. (1954) The life histories of the steelhead rainbow trout (salmo gairdneri gairdneri) and silver salmon (oncorhynchus kisutch) with special reference to waddell creek, California, and recommendations regarding their management. State of California Department of Fish and Game Fish Bulletin 98: 1–375.

66. Williams GC. (1957) Pleiotropy, natural selection, and the evolution of senescence. Evolution 11: 398–411.

67. Pagel MD, Harvey PH. (1988) Recent developments in the analysis of comparative data. Quarterly Review of Biology: 413–40.

68. Garland T, Harvey PH, Ives AR. (1992) Procedures for the analysis of comparative data using phylogenetically independent contrasts. Systematic Biology: 18–32.

69. Luo S, Murphy CT. (2011) Caenorhabditis elegans reproductive aging: Regulation and underlying mechanisms. Genesis 49: 53–65.

70. Reznick D, Bryant M, Holmes D. (2006) The evolution of senescence and post-reproductive lifespan in guppies (poecilia reticulata). PloS Biology 4: e7–e7.

71. Loudon I. (1993) Death in childbirth: An international study of maternal care and maternal mortality 1800–1950. New York, New York: Oxford University Press.

72. Savage-Dunn C, Tokarz R, Wang H, Cohen S, Giannikas C, et al. (2000) SMA-3 smad has specific and critical functions in DBL-l/SMA-6 TGFbeta-related signaling. Developmental Biology 223: 70–6.

73. Robertson L, Mitchell JR. (2013) Benefits of short-term dietary restriction in mammals. Experimental Gerontology In Press.

74. Selesniemi K, Lee H, Tilly JL. (2008) Moderate caloric restriction initiated in rodents during adulthood sustains function of the female reproductive axis into advanced chronological age. Ageing Cell 7: 622–9.

75. Ptak G, Tacconi E, Czernik M, Toschi P, Modlinski J, et al. (2012) Embryonic diapaise is conserved across mammals. PLoS ONE 7: e33027.

76. Grinsted J, Avery B. (1996) A sporadic case of delayed implantation after in-vitro fertilization in the human? Human Reproduction 11: 651–4.

77. Sim C, Denlinger DL. (2008) Insulin signaling and FOXO regulate the overwintering diapause of the mosquitoCu/ex pipiens. Proceedings of the National Academy of Sciences of the United States of America 105: 6777–81.

78. Dickhoff WW. (1989) Salmonids and annual fishes: Death after sex. In: Scanes CJ, Schriebman MP, editors. Development, maturation, and senescence of neuroendocrine systems. New York, NY: Academic, pp. 253–66.

79. Robertson OH. (1961) Prolongation of the life span of kokanee salmon (oncorhynchus nerka kennerlyi) by castration before beginning of gonad development. Proceedings Biological Sciences / The Royal Society 47: 609–21.

80. Berman JR, Kenyon C. (2006) Germ-cell loss extends C. elegans life span through regulation of DAF-16 by kri-1 and lipophilic-hormone signaling. Cell 124: 1055–68.

81. Hsin H, Kenyon C. (1999) Signals from the reproductive system regulate the lifespan of C. elegans. Nature 399: 362–6.

82. Emmerson E, Hardman MJ. (2012) The role of estrogen deficiency in skin ageing and wound healing. Biogerontology 13: 3–20.

83. Aviv A. (2007) Cardiovascular diseases, aging and the gender gap in the human longevity. Journal of the American Society of Hypertension 1: 185–8.

84. Finch CE. (2010) Evolution in health and medicine sackler colloquium: Evolution of the human lifespan and diseases of aging: Roles of infection, inflammation, and nutrition. Proceedings of the National Academy of Sciences of the United States of America 107 Suppl: 1718–24.

85. Kirkwood TB, Rose MR. (1991) Evolution of senescence: Late survival sacrificed for reproduction. Proceedings Biological Sciences / the Royal Society 332: 15–24.

86. Finch CE, Holmes DJ. (2010) Ovarian aging in developmental and evolutionary contexts. Annals of the New York Academy of Sciences 1204: 82–94.

87. Foote AD. (2008) Mortality rate acceleration and post-reproductive lifespan in matrilineal whale species. Biology Letters 4: 189–91.

88. Hendry AP, Morbey YE, Berg OK, Wenburg JK. (2004) Adaptive variation in senescence: Reproductive lifespan in a wild salmon population. Proceedings Biological Sciences / The Royal Society 271: 259–66.

